# Pleiotropy between neuroticism and physical and mental health: findings from 108 038 men and women in UK Biobank

**DOI:** 10.1101/031138

**Authors:** Catharine R Gale, Saskia P Hagenaars, Gail Davies, W David Hill, David CM Liewald, Breda Cullen, International Consortium for Blood Pressure GWAS, CHARGE consortium Aging and Longevity Group, Jill Pell, Andrew M McIntosh, Daniel J Smith, Ian J Deary, Sarah E Harris

**Affiliations:** Centre for Cognitive Ageing and Cognitive Epidemiology, University of Edinburgh, 7 George Square, Edinburgh EH8 9JZ, UK; Department of Psychology, University of Edinburgh, 7 George Square, Edinburgh, EH8 9JZ, UK; MRC Lifecourse Epidemiology Unit, University of Southampton, Southampton, SO16 6YD, UK; Division of Psychiatry, University of Edinburgh, Edinburgh, EH10 5HF, UK; Institute of Health and Wellbeing, University of Glasgow, Glasgow, G12 8RZ, UK; Medical Genetics Section, University of Edinburgh Centre for Genomic and Experimental Medicine and MRC Institute of Genetics and Molecular Medicine, Western General Hospital, Crewe Road, Edinburgh EH4 2XU, UK

## Abstract

There is considerable evidence that people with higher levels of the personality trait neuroticism have an increased risk of several types of mental disorder. Higher neuroticism has also been associated, less consistently, with increased risk of various physical health outcomes. We hypothesised that these associations may, in part, be due to shared genetic influences. We tested for pleiotropy between neuroticism and 12 mental and physical diseases or health traits using linkage disequilibrium regression and polygenic profile scoring. Genetic correlations were derived between neuroticism scores in 108 038 people in UK Biobank and health-related measures from 12 large genome-wide association studies (GWAS). Summary information for the 12 GWAS was used to create polygenic risk scores for the health-related measures in the UK Biobank participants. Associations between the health-related polygenic scores and neuroticism were examined using regression, adjusting for age, sex, genotyping batch, genotyping array, assessment centre, and population stratification. Genetic correlations were identified between neuroticism and anorexia nervosa (r_g_ = 0.17), major depressive disorder (r_g_ = 0.66) and schizophrenia (r_g_ = 0.21). Polygenic risk for several health-related measures were associated with neuroticism, in a positive direction in the case of bipolar disorder (β = 0.017), major depressive disorder (β = 0.036), schizophrenia (β = 0.036), and coronary artery disease (β = 0.011), and in a negative direction in the case of BMI (β = −0.0095). These findings indicate that a high level of pleiotropy exists between neuroticism and some measures of mental and physical health, particularly major depressive disorder and schizophrenia.

## Introduction

There is considerable evidence that the personality trait of neuroticism^1^—which describes stable individual differences in the tendency to experience negative emotions—has profound significance for public health.^2^ People who are higher in neuroticism have an increased risk of developing Axis I psychopathology, especially the common mental disorders such as mood, anxiety, somatoform and substance use disorders, and also schizophrenia, bipolar disorder and attention deficit hyperactivity disorder (ADHD). ^3-6^ Higher neuroticism is associated with increased likelihood of diagnosis with Axis II personality disorders^7^ and with greater comorbidity between internalising disorders (such as major depression, generalised anxiety, panic disorders and phobias) and externalising disorders (such as alcohol and drug dependence, antisocial personality and conduct disorders).^8^ There is evidence that higher neuroticism is linked with risk of developing Alzheimer’s disease.^9^ People who are higher in neuroticism tend to make greater use of mental health services, regardless of whether they have a mental disorder,^10^ perhaps because they are more likely to perceive a need for care.^11^ The estimated economic costs of neuroticism in terms of health care use and absenteeism are massive.^12^ Much of these costs relate to reported chronic somatic conditions.^12^ Higher neuroticism has been linked with increased somatic complaints,^13, 14^ with perception of poorer health,^15, 16^ with future somatic multi-morbidity, as assessed by a count of reported chronic conditions,^17^ and with increased likelihood of reporting a range of physical health problems.^18^

Evidence that neuroticism is predictive of objectively-assessed physical health is still relatively sparse and findings to date are often mixed. For example, whereas some prospective studies have found that higher neuroticism increases mortality from all causes^19^ or coronary heart disease,^20^ and is predictive of raised blood pressure^21^ or body mass index (BMI),^22^ others have found no such association.^23-26^ In a pooled analysis of data from five cohorts, there was no consistent association between neuroticism and incidence of type 2 diabetes: higher neuroticism was linked with increased risk in one cohort, but not in the others.^27^

Part of the explanation for associations between neuroticism and these various mental and physical health outcomes may be due to shared genetic influences. Twin and adoption studies suggest that genetic influences account for between a third and a half of individual differences in neuroticism.^1^ Many physical and mental illnesses and health-related measures also show moderate heritability.^28^ Twin studies have shown that there is considerable overlap between the genetic factors that influence variations in neuroticism and those that determine risk of depression and other internalizing disorders.^29, 30^ It is now possible to test for such pleiotropy in associations between neuroticism and health outcomes using data from single nucleotide polymorphism (SNP) genotyping in unrelated individuals, making it possible to carry out much larger studies without the assumptions made by twin-study methods. A recent genome-wide association meta-analysis based on data from over 70 000 individuals found that neuroticism is influenced by many genetic variants of small effect, i.e. a polygenic effect, that also influence risk of major depressive disorder.^31^ Whether there is pleiotropy between neuroticism and other mental disorders or with physical health outcomes is unclear.

Testing for pleiotropy using SNP-based genetic data can be carried out in several ways. Linkage disequilibrium (LD) regression calculates genetic correlations between health measures using the summary results of genome-wide association studies (GWAS).^32^ Polygenic risk scoring^33^ uses summary GWAS data for a given illness or health trait to test whether polygenic liability to that illness/trait is associated with phenotypes for that illness/trait (e.g. neuroticism scores) or others measured in an independent sample. Polygenic risk of neuroticism was recently associated with major depressive disorder.^31^

In the present study we aimed to discover whether shared genetic aetiology explains part of the associations between neuroticism and various physical and mental health outcomes. We used data on over 108 000 UK Biobank participants who completed a questionnaire on neuroticism and provided DNA for genome-wide genotyping. Using summary data from GWAS meta-analyses on 12 health-related measures, we tested for neuroticism-health pleiotropy using two complementary methods. First, we used LD regression to derive genetic correlations between health-related measures and neuroticism. Second, we calculated the associations between polygenic risk scores for the 12 health-related measures and the neuroticism phenotype in UK Biobank participants.

## Methods

### Participants

The participants in this study took part in the baseline survey of UK Biobank. ^34^ (http://www.ukbiobank.ac.uk). UK Biobank was set up as a resource for identifying determinants of disease in middle aged and older people. Between 2006 and 2010, 502 655 community-dwelling people aged between 37 and 73 years and living in the United Kingdom were recruited to the study. They underwent assessments of cognitive and physical functions, mood and personality. They provided blood, urine, and saliva samples for future analysis, completed questionnaires about their social backgrounds and lifestyle, and agreed to have their health followed longitudinally. UK Biobank received ethical approval from the Research Ethics Committee (REC reference 11/NW/0382).

For the present study, genome-wide genotyping data were available on 112 151 individuals (58 914 female) aged 40-73 years (mean age = 56.9 years, SD = 7.9) after the quality control process (see below).

## Procedures

### Neuroticism

Participants completed the Neuroticism scale of the Eysenck Personality Questionnaire-Revised Short Form (EPQ-R Short Form).^35^ This scale has been concurrently validated in older people against two of the most widely-used measures of neuroticism, taken from the International Personality Item Pool (IPIP) and the NEO-Five Factor Inventory (NEO-FFI); it correlated −0.84 with the IPIP-Emotional Stability scale and 0.85 with the NEO-FFI Neuroticism scale.^36^ A previous study found a high genetic correlation (0.91) between the EPQ-R Short Form Neuroticism scale and psychological distress assessed in a non-psychiatric population using the 30-item General Health Questionnaire.^37^

### Genotyping and quality control

152 729 UK Biobank blood samples were genotyped using either the UK BiLEVE^38^ array (N = 49 979) or the UK Biobank axiom array (N = 102 750). A full description of the genotyping process is available in the Supplementary Materials. Quality control was performed by Affymetrix, the Wellcome Trust Centre for Human Genetics, and by the present authors; this included removal of participants based on missingness, relatedness, gender mismatch, non-British ancestry, and other criteria, and is described in the Supplementary Materials.

### Genome-wide association analyses (GWAS) in the UK Biobank sample

Genome-wide association analyses were performed on the neuroticism measure in order to use the summary results for linkage disequilibrium regression. Details of the GWAS procedures are provided in the Supplementary Materials.

### Curation of summary results from GWAS consortia on health-related variables

In order to conduct LD regression and polygenic profile score analyses between the UK Biobank neuroticism data and the genetic predisposition to mental and physical health outcomes, we gathered summary data from published meta-analyses on 12 health-related measures: six relating to mental health (ADHD, Alzheimer’s disease, anorexia nervosa, bipolar disorder, major depressive disorder and schizophrenia) and six relating to physical health (systolic and diastolic blood pressure, BMI, coronary artery disease, longevity, and type 2 diabetes). Details of these health-related variables, the consortia’s websites, key references, and number of subjects included in each consortium’s genome-wide association study are given in Supplementary Table 1.

**Table 1.** Genetic correlations between neuroticism documented in UK Biobank and mental and physical health-related traits curated from GWAS consortia. Statistically significant p-values (after False Discovery Rate correction-p<2.39×10^−5^) are shown in bold.

| Trait Category | Traits from GWAS consortia | Neuroticism (n = 108,038) |  |  |
| --- | --- | --- | --- | --- |
| | | $r_g$ | SE | p |
| **Mental health** | ADHD | 0.060 | 0.082 | 0.4681 |
|  | Alzheimer's Disease | 0.118 | 0.080 | 0.1378 |
|  | Alzheimer's Disease (500kb) | 0.091 | 0.063 | 0.1514 |
|  | Anorexia Nervosa | 0.174 | 0.04 | **$2.36 \times 10^{-5}$** |
|  | Bipolar Disorder | 0.083 | 0.052 | 0.1096 |
|  | Major Depressive Disorder | 0.659 | 0.087 | **$2.75 \times 10^{-14}$** |
|  | Schizophrenia | 0.212 | 0.050 | **$2.39 \times 10^{-5}$** |
| **Physical health** | Blood Pressure: Diastolic | -0.011 | 0.078 | 0.887 |
|  | Blood Pressure: Systolic | -0.051 | 0.060 | 0.3959 |
|  | BMI | -0.036 | 0.028 | 0.1921 |
|  | Coronary Artery Disease | 0.104 | 0.053 | 0.0489 |
|  | Longevity | -0.083 | 0.081 | 0.3011 |
|  | Type 2 Diabetes | -0.145 | 0.07 | 0.0389 |

## Statistical Analysis

### Computing genetic associations between neuroticism and health-related variables

We use two methods to compute genetic associations between neuroticism and the health-related variables, LD regression and polygenic profile/risk scoring. Each provides a different metric to infer the existence of pleiotropy between pairs of traits. LD regression was used to derive genetic correlations to determine the degree to which the polygenic architecture of a trait overlaps with that of another. The polygenic risk score method was used to test the extent to which the polygenic information from GWASs of health-related variables could predict neuroticism in the UK Biobank participants. Both LD regression and polygenic risk scores are dependent on the traits analysed being highly polygenic in nature, i.e. where a large number of variants of small effect contribute toward phenotypic variation.^32, 39^

### LD regression

In order to quantify the extent of pleiotropy between neuroticism, measured in UK Biobank, and the collated health traits, we used LD regression.^32, 40^ This is a class of techniques that exploits the correlational structure of the single nucleotide polymorphisms (SNPs) found across the genome (we provide more details of LD regression in Supplementary material). Here, we use LD regression to derive genetic correlations between neuroticism and health-related measures using 12 large GWAS consortia data sets that enable pleiotropy of their health-related traits to be quantified with the neuroticism trait in UK Biobank. We followed the data processing pipeline devised by Bulik-Sullivan et al.^32, 40^ described in more detail in the Supplementary Materials. In order to ensure that the genetic correlation for the Alzheimer’s disease phenotype was not driven by a single locus or biased the fit of the regression model, a 500kb region centred on the *APOE* locus was removed and this phenotype was re-run. This additional model is referred to in the Tables below as ‘Alzheimer’s disease (500kb)’.

### Polygenic profiling

The UK Biobank genotyping data required recoding from numeric (1, 2) allele coding to standard ACGT format prior to being used in polygenic profile scoring analyses. This was achieved using a bespoke programme developed by one of the present authors (DCML), details of which are provided in the Supplementary Materials.

Polygenic profiles were created for 12 health-related phenotypes (see Table 3, Supplementary Table 2) in all genotyped participants using PRSice. ^41^ This software calculates the sum of alleles associated with the phenotype of interest across many genetic loci, weighted by their effect sizes estimated from a GWAS of that phenotype in an independent sample. Prior to creating the scores, SNPs with a minor allele frequency < 0.01 were removed and clumping was used to obtain SNPs in linkage equilibrium with an r^2^ < 0.25 within a 200bp window. Multiple scores were then created for each phenotype containing SNPs selected according to the significance of their association with the phenotype. The GWAS summary data for the 12 health-related phenotypes were used to create five polygenic profiles for each in the UK Biobank participants, at thresholds of p < 0·01, p < 0·05, p < 0·1, p < 0·5 and all SNPs. The most predictive threshold will be presented in the main tables of the paper. The full results, including all five thresholds, can be found in the Supplementary Table 2.

Associations between the polygenic profiles and neuroticism were examined in linear regression models, adjusting for age at measurement, sex, genotyping batch and array, assessment centre, and the first ten genetic principal components to adjust for population stratification. We corrected for multiple testing across all polygenic profile scores at all significance thresholds for associations with neuroticism using the False Discovery Rate (FDR) method.^42^

## Results

Within UK Biobank, 108 038 individuals with genotype data completed the Neuroticism scale of the Eysenck Personality Questionnaire-Revised Short Form. Their mean (SD) score for neuroticism was 4.02 (3.17).

Table 1 shows the genetic correlations obtained using LD regression between neuroticism in UK Biobank and the published GWAS results on the health-related traits. Neuroticism showed significant positive genetic correlations with three traits, all related to mental health, namely major depressive disorder (r_g_ = 0.66), schizophrenia (r_g_ = 0.21) and anorexia nervosa (r_g_ = 0.17). There were no significant genetic correlations between neuroticism and either the other mental health-related traits [attention-deficit hyperactivity disorder (ADHD), Alzheimer’s disease and bipolar disorder] or any of the physical health-related traits (systolic and diastolic blood pressure, BMI, coronary artery disease, type 2 diabetes or longevity. These results are shown graphically in Figure 1.

**Figure 1.**
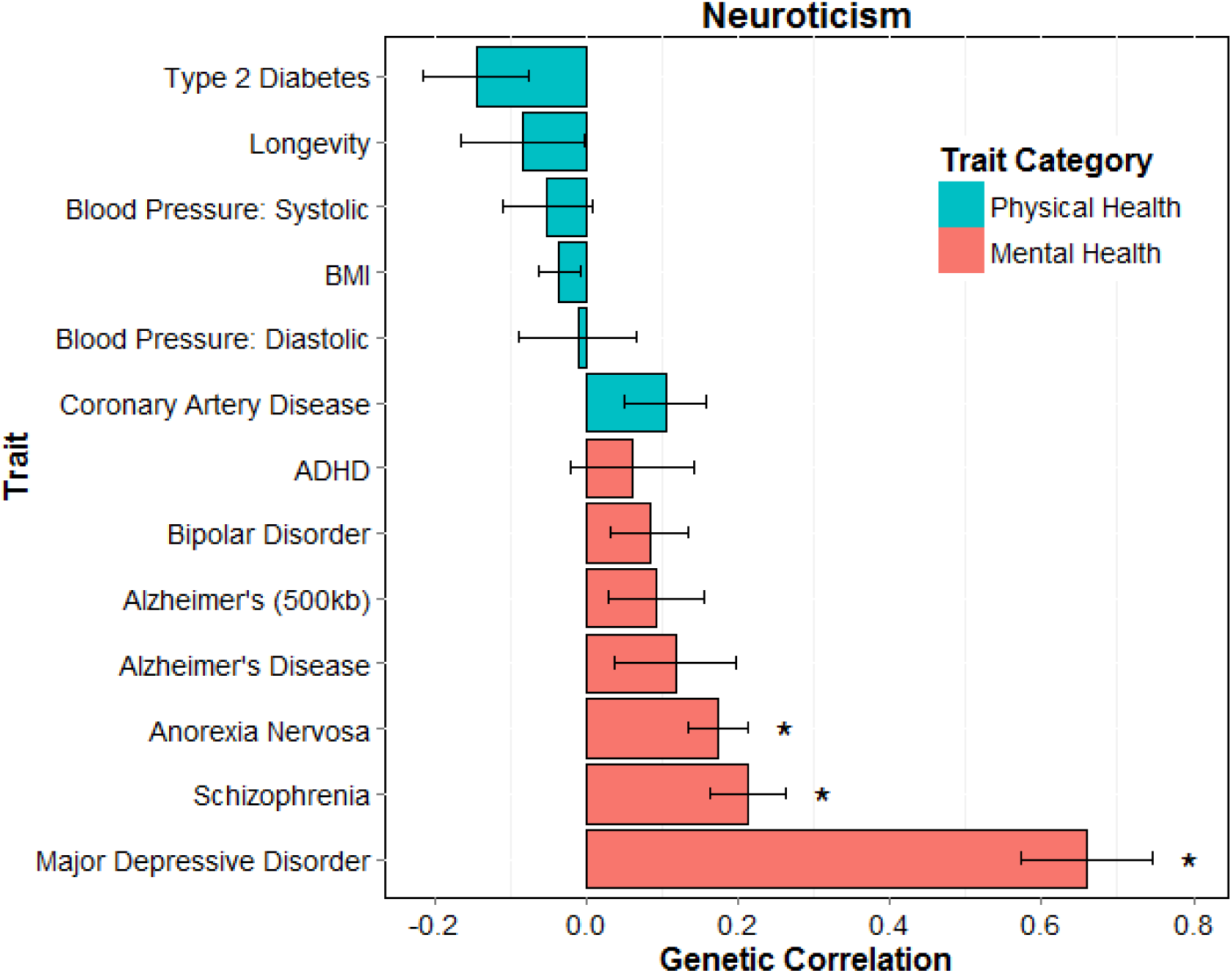
Barplot of genetic correlations (SE) calculated using LD regression between neuroticism in UK Biobank and mental and physical health measures from GWAS consortia.*, p<2.39×10−5

Table 2 shows the results of the polygenic risk scoring, using the most predictive threshold of the five that were created. Higher polygenic risk for three mental health-related traits, bipolar disorder, major depressive disorder, and schizophrenia, was significantly associated with higher levels of neuroticism (standardised β between 0.017 – 0.036). There were no significant associations between neuroticism and polygenic risk for the other mental health-related traits examined, namely ADHD, Alzheimer’s disease, and anorexia nervosa. Polygenic risk scores for two physical health-related traits were significantly associated with neuroticism: higher polygenic risk for BMI was associated with lower levels of neuroticism (β = −0.0095, *p* = 0.0015), and higher polygenic risk for coronary artery disease was associated with higher levels of neuroticism (β = 0.011, *p* = 0.0003). No significant associations were found between polygenic risk for the other physical health-related traits (systolic and diastolic blood pressure, type 2 diabetes or longevity) and neuroticism. To test whether the FDR significant association between polygenic risk for coronary artery disease and neuroticism was confounded by individuals diagnosed with cardiovascular disease, 2717 individuals who had had a heart attack and 2468 individuals with angina were removed from the regression analysis. Our estimate of the association between polygenic risk of coronary artery disease and neuroticism was unchanged by this exclusion. The complete polygenic risk score results, including all five thresholds, are shown in Supplementary Table 2.

**Table 2.** Associations between polygenic risk scores for mental and physical health-related traits created from GWAS consortia summary data and neuroticism in UK Biobank participants, adjusted for age, sex, assessment centre, genotyping batch and array, and ten principal components for population stratification. The associations between the polygenic risk scores and neuroticism with the largest effect size (threshold) are presented. Statistically significant p-values (after False Discovery Rate correction-p<0.0061) are shown in bold.

| Trait Category | Traits from GWAS consortia | Neuroticism (n = 108,038) |  |  |
| --- | --- | --- | --- | --- |
| | | Threshold | $\beta$ | p |
| **Mental health** | ADHD | 0.05 | 0.0045 | 0.1292 |
|  | Alzheimer's Disease | 0.05 | -0.0065 | 0.0312 |
|  | Anorexia Nervosa | 0.1 | 0.0054 | 0.0735 |
|  | Bipolar Disorder | 0.5 | 0.0171 | **$1.71 \times 10^{-8}$** |
|  | Major Depressive Disorder | 1 | 0.0357 | **$1.23 \times 10^{-31}$** |
|  | Schizophrenia | 0.1 | 0.0359 | **$7.88 \times 10^{-32}$** |
| **Physical health** | Blood Pressure: Diastolic | 0.1 | -0.0047 | 0.1167 |
|  | Blood Pressure: Systolic | 0.05 | -0.0016 | 0.5969 |
|  | BMI | 0.01 | -0.0095 | **0.0015** |
|  | Coronary Artery Disease | 0.1 | 0.0109 | **0.0003** |
|  | Longevity | 0.05 | -0.0051 | 0.0917 |
|  | Type 2 Diabetes | 0.01 | -0.0031 | 0.2989 |

## Discussion

In this study of 108 038 men and women from UK Biobank who had been genotyped and assessed for neuroticism, we exploited the summary results of 12 large international GWAS consortia to examine whether there is pleiotropy between neuroticism and a range of physical and mental health outcomes using two methods, LD regression and polygenic profile scoring. As regards the six mental health outcomes, we found consistent evidence of pleiotropy between neuroticism and both major depression and schizophrenia with these methods, showing that, to a significant degree, the same genetic variants are responsible for the heritability of each pair of phenotypes and that genetic variants associated with major depression or schizophrenia in GWAS consortia are significantly predictive of variation in neuroticism in UK Biobank. There was some evidence for pleiotropy between neuroticism and both bipolar disorder and anorexia nervosa on the basis of results from polygenic profile scoring or LD regression respectively, but the extent of these associations varied according to the method used. We found no evidence of pleiotropy between neuroticism and the other mental health outcomes examined—ADHD and Alzheimer’s disease. Of the six physical health outcomes studied, none showed evidence of pleiotropy with neuroticism on the basis of the genetic correlations obtained from LD regression, but there was some indication of genetic overlap between neuroticism and both coronary artery disease and BMI. Higher polygenic risk for coronary artery disease was significantly associated with higher levels of neuroticism and polygenic risk of higher BMI was associated with lower levels of neuroticism. No associations were found between polygenic risk for the other physical health-related traits (systolic and diastolic blood pressure, type 2 diabetes or longevity) and neuroticism.

Previous investigations into pleiotropy between neuroticism and mental health outcomes using polygenic risk profiling have found evidence of substantial shared genetic aetiology between neuroticism and major depression.^31, 43, 44^ Our observations in the present, much larger, sample confirm those findings and for the first time quantify the extent to which the same genetic variants are responsible for the heritability in these two phenotypes, using an additional metric, LD regression.^32^ The relatively high genetic correlation between neuroticism and major depression (r_g_ = 0.66), identified in our study, is similar to the genetic correlation identified in a previous twin study (r_g_ = 0.43).^30^ Our results also provide the first evidence, to our knowledge, that the phenotypic correlations found between neuroticism and schizophrenia^45, 46^ are due at least in part to genetic overlap. The extent of pleiotropy between anorexia nervosa or bipolar disorder and neuroticism has been unclear, though there is some, though limited, evidence to link both disorders phenotypically with neuroticism.^5, 47^ A previous study found no significant association between polygenic risk scores for neuroticism and bipolar disorder, but the sample size was small.^44^ In this much larger sample, higher polygenic risk scores for bipolar disorder were significantly predictive of higher neuroticism, but results of LD regression showed little indication of genetic correlation between neuroticism and this disorder. We found a small but highly significant genetic correlation between neuroticism and anorexia (r_g_ = 0.17), but there was no association between polygenic risk scores for this condition and neuroticism. Although there is now considerable evidence that neuroticism is a risk factor for the development of Alzheimer’s disease,^9^ there was no indication in our analyses that shared genes account for this link.

So far as we are aware, there have been no previous investigations of the extent of pleiotropy between neuroticism and physical health outcomes. This may be because, whereas there is considerable evidence for phenotypic associations between higher neuroticism and poorer self-rated health or greater somatic complaints, ^13-18^ fewer studies have examined neuroticism as a predictor of objectively-measured physical health outcomes and findings on such outcomes as coronary heart disease, blood pressure, BMI and all-cause mortality have been inconsistent. ^19-26^ Of the six objectively measured physical health outcomes included in the current study, two showed evidence of a degree of genetic pleiotropy with neuroticism: coronary artery disease and BMI. Neither demonstrated any measurable genetic correlation with neuroticism, but higher polygenic risk score for coronary artery disease was associated with higher neuroticism. This is consistent with the finding in pooled data from three cohorts that higher neuroticism was associated with increased mortality from coronary heart disease.^20^ Higher polygenic risk for BMI was associated with lower neuroticism. The direction of this association was unexpected. Findings on the phenotypic relationships between neuroticism and BMI have produced inconsistent results: one study showed higher neuroticism was associated with higher BMI,^22^ but another found no association.^25^

The chief strength of our study is the large sample size which permits powerful, robust tests of genetic association. Secondly, all the UK Biobank genetic data were processed at the same location and on the same platform. Finally, use of summary data from 12 large international GWAS consortia studies allowed us to perform a comprehensive examination of the degree of pleiotropy between neuroticism and a range of physical and mental health-related phenotypes, and to produce many of the first estimates of the genetic correlation between neuroticism and these phenotypes.

Our study also has some limitations. Firstly, the GWAS studies we curated to carry out LD regression and extract polygenic risk scores often involved meta-analyses of results from datasets with considerable heterogeneity in sample size, genome-wide imputation quality and measurement of phenotypes. With larger and more consistent independent datasets, it should be possible to use the polygenic risk scores to predict more variance in neuroticism. Secondly, we restricted the genotyped samples to individuals of white British ancestry in order to minimise any influence of population structure. Our results therefore need to be replicated in large samples with different genetic backgrounds as we did not have the power to model data from UK Biobank individuals of other ancestries.

The polygenic risk score most predictive threshold varied between health traits, suggesting that the amount of shared genetic aetiology between neuroticism and each of the health traits differs. Although testing multiple thresholds may be deemed to increase the multiple testing problem, it should be noted that the SNPs included in each threshold are not independent. Also, at least for major depressive disorder and schizophrenia, pleiotropy was quantified using LD regression, which involved only a single test per pair of phenotypes. The estimate of neuroticism variance explained by each polygenic profile should be considered as the minimum estimate of the variance explained. Due to pruning SNPs in LD, the polygenic risk score method makes the assumption of a single causal variant being tagged in each LD block considered. If this assumption is not true for the phenotypes considered, the proportion of variance explained will be underestimated here.

In this large sample from UK Biobank, we aimed to discover whether shared genetic aetiology explained part of the associations between neuroticism and various physical and mental health outcomes. Our findings suggest that associations between neuroticism and several mental health outcomes including major depression, schizophrenia, bipolar disorder and anorexia nervosa are in part due to shared genetic influences. We found that polygenic risk scores for coronary artery disease and BMI were predictive of neuroticism scores. This large-scale mapping of the extent of pleiotropy between neuroticism and physical and mental health outcomes adds to our understanding of the cause of links between this important personality trait and later health.

## Acknowledgments

This work was supported by the Biotechnology and Biological Sciences Research Council and Medical Research Council (MR/K026992/1), Age UK’s Disconnected Mind Project, Wellcome Trust (104036/Z/14/Z).

## References

1. Matthews G, Deary IJ, Whiteman MC. Personality traits. 3rd edn. Cambridge University Press: Cambridge, 2009.

2. Lahey BB. Public health significance of neuroticism. Am Psychol 2009; 64(4): 241–256.

3. Kotov R, Gamez W, Schmidt F, Watson D. Linking “big” personality traits to anxiety, depressive, and substance use disorders: a meta-analysis. Psychol Bull 2010; 136(5): 768–821.

4. Malouff JM, Thorsteinsson EB, Schutte NS. The relationship between the five-factor model of personality and symptoms of clinical disorders: a meta-analysis. Journal of Psychopathology & Behavioral Assessment 2005; 27: 101–114.

5. Lonnqvist JE, Verkasalo M, Haukka J, Nyman K, Tiihonen J, Laaksonen I et al. Premorbid personality factors in schizophrenia and bipolar disorder: results from a large cohort study of male conscripts. J Abnorm Psychol 2009; 118(2): 418–423.

6. Michielsen M, Comijs HC, Semeijn EJ, Beekman AT, Deeg DJ, Kooij JJ. Attention deficit hyperactivity disorder and personality characteristics in older adults in the general Dutch population. Am J Geriatr Psychiatry 2014; 22(12): 1623–1632.

7. Saulsman LM, Page AC. The five-factor model and personality disorder empirical literature: A meta-analytic review. Clin Psychol Rev 2004; 23(8): 1055–1085.

8. Khan AA, Jacobson KC, Gardner CO, Prescott CA, Kendler KS. Personality and comorbidity of common psychiatric disorders. Br J Psychiatry 2005; 186: 190–196.

9. Terracciano A, Sutin AR, An Y, O’Brien RJ, Ferrucci L, Zonderman AB et al. Personality and risk of Alzheimer’s disease: new data and meta-analysis. Alzheimers Dement 2014; 10(2): 179–186.

10. ten Have M, Oldehinkel A, Vollebergh W, Ormel J. Does neuroticism explain variations in care service use for mental health problems in the general population? Results from the Netherlands Mental Health Survey and Incidence Study (NEMESIS). Soc Psychiatry Psychiatr Epidemiol 2005; 40(6): 425–431.

11. Seekles WM, Cuijpers P, van de Ven P, Penninx BW, Verhaak PF, Beekman AT et al. Personality and perceived need for mental health care among primary care patients. J Affect Disord 2012; 136(3): 666–674.

12. Cuijpers P, Smit F, Penninx BW, de Graaf R, ten Have M, Beekman AT. Economic costs of neuroticism: a population-based study. Arch Gen Psychiatry 2010; 67(10): 1086–1093.

13. Costa PT, Mccrae RR. Neuroticism, Somatic Complaints, and Disease - Is the Bark Worse Than the Bite. Journal of Personality 1987; 55(2): 299–316.

14. Jorm AF, Christensen H, Henderson S, Korten AE, Mackinnon AJ, Scott R. Neuroticism and Self-Reported Health in an Elderly Community Sample. Personality and Individual Differences 1993; 15(5): 515–521.

15. Goodwin R, Engstrom G. Personality and the perception of health in the general population. Psychol Med 2002; 32(2): 325–332.

16. Watson D, Pennebaker JW. Health complaints, stress, and distress: exploring the central role of negative affectivity. Psychol Rev 1989; 96(2): 234–254.

17. Neeleman J, Bijl R, Ormel J. Neuroticism, a central link between somatic and psychiatric morbidity: path analysis of prospective data. Psychol Med 2004; 34(3): 521–531.

18. Goodwin RD, Cox BJ, Clara I. Neuroticism and physical disorders among adults in the community: Results from the national comorbidity survey. Journal of Behavioral Medicine 2006; 29(3): 229–238.

19. Shipley BA, Weiss A, Der G, Taylor MD, Deary IJ. Neuroticism, extraversion, and mortality in the UK Health and Lifestyle Survey: a 21-year prospective cohort study. Psychosom Med 2007; 69(9): 923–931.

20. Jokela M, Pulkki-Raback L, Elovainio M, Kivimaki M. Personality traits as risk factors for stroke and coronary heart disease mortality: pooled analysis of three cohort studies. Journal of Behavioral Medicine 2014; 37: 881–889.

21. Turiano NA, Pitzer L, Armour C, Karlamangla A, Ryff CD, Mroczek DK. Personality trait level and change as predictors of health outcomes: findings from a national study of Americans (MIDUS). J Gerontol B Psychol Sci Soc Sci 2012; 67(1): 4–12.

22. Magee CA, Heaven PCL. Big-Five personality factors, obesity and 2-year weight gain in Australian adults. J Res Pers 2011; 45(3): 332–335.

23. Nakaya N, Tsubono Y, Hosokawa T, Hozawa A, Kuriyama S, Fukudo S et al. Personality and mortality from ischemic heart disease and stroke. Clin Exp Hypertens 2005; 27(2–3): 297–305.

24. Jokela M, Batty GD, Nyberg ST, Virtanen M, Nabi H, Singh-Manoux A et al. Personality and all-cause mortality: individual-participant meta-analysis of 3,947 deaths in 76,150 adults. Am J Epidemiol 2013; 178(5): 667–675.

25. Mottus R, McNeill G, Jia X, Craig LC, Starr JM, Deary IJ. The associations between personality, diet and body mass index in older people. Health Psychol 2013; 32(4): 353–360.

26. Almada SJ, Zonderman AB, Shekelle RB, Dyer AR, Daviglus ML, Costa PT, Jr. et al. Neuroticism and cynicism and risk of death in middle-aged men: the Western Electric Study. Psychosom Med 1991; 53(2): 165–175.

27. Jokela M, Elovainio M, Nyberg ST, Tabak AG, Hintsa T, Batty GD et al. Personality and risk of diabetes in adults: pooled analysis of 5 cohort studies. Health Psychol 2014; 33: 1618–1618.

28. Polderman TJ, Benyamin B, de Leeuw CA, Sullivan PF, van Bochoven A, Visscher PM et al. Meta-analysis of the heritability of human traits based on fifty years of twin studies. Nature genetics 2015; 47(7): 702–709.

29. Hettema JM, Neale MC, Myers JM, Prescott CA, Kendler KS. A population-based twin study of the relationship between neuroticism and internalizing disorders. Am J Psychiatry 2006; 163(5): 857–864.

30. Kendler KS, Myers J. The genetic and environmental relationship between major depression and the five-factor model of personality. Psychol Med 2010; 40(5): 801–806.

31. Genetics of Personality C, de Moor MH, van den Berg SM, Verweij KJ, Krueger RF, Luciano M et al. Meta-analysis of Genome-wide Association Studies for Neuroticism, and the Polygenic Association With Major Depressive Disorder. JAMA Psychiatry 2015; 72(7): 642–650.

32. Bulik-Sullivan BK, Loh PR, Finucane HK, Ripke S, Yang J, Schizophrenia Working Group of the Psychiatric Genomics C et al. LD Score regression distinguishes confounding from polygenicity in genome-wide association studies. Nature genetics 2015; 47(3): 291–295.

33. Purcell S, Neale B, Todd-Brown K, Thomas L, Ferreira MA, Bender D et al. PLINK: a tool set for whole-genome association and population-based linkage analyses. Am J Hum Genet 2007; 81(3): 559–575.

34. Sudlow C, Gallacher J, Allen N, Beral V, Burton P, Danesh J et al. UK biobank: an open access resource for identifying the causes of a wide range of complex diseases of middle and old age. PLoS Med 2015; 12(3): e1001779.

35. Deary IJ, Bedford A. Some origins and evolution of the EPQ-R (short form) Neuroticism and Extraversion items. Personality and Individual Differences 2011; 50(8): 1213–1217.

36. Gow AJ, Whiteman MC, Pattie A, Deary IJ. Goldberg’s ‘IPIP’ Big-Five factor markers: Internal consistency and concurrent validation in Scotland. Personality and Individual Differences 2005; 39(2): 317–329.

37. Ivkovic V, Vitart V, Rudan I, Janicijevic B, Smolej-Narancic N, Skaric-Juric T et al. The Eysenck personality factors: Psychometric reliability, heritability and phenotypic and genetic correlations with psychological distress in an isolated Croatian population. Personality and Individual Differences 2007; 42: 123–133.

38. Wain LV, Shrine N, Miller S, et al. Novel insights into the genetics of smoking behaviour, lung function, and chronic obstructive pulmonary disease (UK BiLEVE): a genetic association study in UK Biobank. The Lancet Respiratory Medicine 2015 doi:10.1016/S2213–2600(15)00283–0.

39. International Schizophrenia C, Purcell SM, Wray NR, Stone JL, Visscher PM, O’Donovan MC et al. Common polygenic variation contributes to risk of schizophrenia and bipolar disorder. Nature 2009; 460(7256): 748–752.

40. Bulik-Sullivan B, Finucane HK, Anttila V, Gusev A, Day FR, Perry JR et al. An Atlas of Genetic Correlations across Human Diseases and Traits. bioRxiv 2015; doi: http://dx.doi.org/10.1101/014498

41. Eusden J, Lewis CM, O’Reilly PF. PRSice: Polygenic Risk Score software. Bioinformatics 2015; 31: 1466–1468.

42. Benjamini Y, Hochberg Y. Controlling the False Discovery Rate - a Practical and Powerful Approach to Multiple Testing. J Roy Stat Soc B Met 1995; 57(1): 289–300.

43. Luciano M, Huffman JE, Arias-Vasquez A, Vinkhuyzen AA, Middeldorp CM, Giegling I et al. Genome-wide association uncovers shared genetic effects among personality traits and mood states. American journal of medical genetics Part B, Neuropsychiatric genetics: the official publication of the International Society of Psychiatric Genetics 2012; 159B(6): 684–695.

44. Middeldorp CM, de Moor MH, McGrath LM, Gordon SD, Blackwood DH, Costa PT et al. The genetic association between personality and major depression or bipolar disorder. A polygenic score analysis using genome-wide association data. Translational psychiatry 2011; 1: e50.

45. Horan WP, Blanchard JJ, Clark LA, Green MF. Affective traits in schizophrenia and schizotypy. Schizophr Bull 2008; 34(5): 856–874.

46. Van Os J, Jones PB. Neuroticism as a risk factor for schizophrenia. Psychol Med 2001; 31(6): 1129–1134.

47. Bulik CM, Sullivan PF, Tozzi F, Furberg H, Lichtenstein P, Pedersen NL. Prevalence, heritability, and prospective risk factors for anorexia nervosa. Arch Gen Psychiatry 2006; 63(3): 305–312.

